## Supplementary Information compiled pdf containg 15 SI Figures, 3 SI Notes, SI Movie legends, and SI references for "Correlating fluorescence microscopy, optical and magnetic tweezers to study single chiral biopolymers such as DNA"

Supplementary Information contains:

**Supplementary Notes 1-3**

**Supplementary Figures 1-15**

**Supplementary Movie Legends 1-16**

**Supplementary Note 1: Evaluation of the magnetic force to show limited B-field influence on the tether**

The B field in the sample space due to the magnetic tweezers is numerically evaluated using the Biot-Savart Law:

$$d\boldsymbol{B} = \frac{\mu_0}{4\pi} \cdot \frac{I \cdot d\boldsymbol{s} \cdot \sin(\theta)}{\boldsymbol{r}^2}$$

Where $\mu_0$ is the permeability of free space, which is equal to $4\pi \times 10^{-7}$ $TmA^{-1}$, $I$ is the amplitude of the current flowing in the coils, $d\boldsymbol{s}$ is the length of a small element of the coil, $r$ is the position vector of the point in question drawn from the current element and $\theta$ is the angle between the two. The B field is obtained by integrating $d\boldsymbol{B}$ over all the coils.

$$\boldsymbol{B}(\boldsymbol{r}) = \int_{coil\ start}^{coil\ end} d\boldsymbol{B}$$

The near Helmoholtz design of the coils makes the B field as uniform as possible to minimize the B force potentially applied on the magnetic bead. We found from the above evaluation that when the pair of smaller coils alone are turned on, the B field changes most with distance. But even in this case the B field strength increases only 0.002% over a distance of 40 µm from the centre of the trap. See Supplementary Fig. 1.

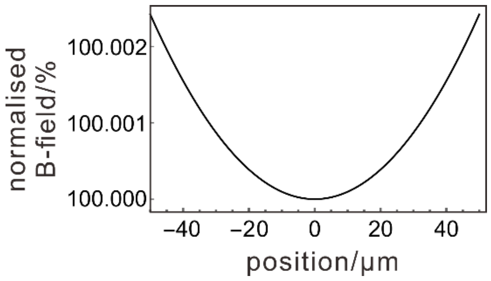


**Supplementary Fig. 1. COMBI-Tweez B field is highly uniform in vicinity of the sample.** Normalised B field over a region of 80 µm around the centre of the magnetic trap when the pair of smaller coils are turned on.

The magnetic force, $\boldsymbol{F}$, is the change of B field with respect to space:

$$\boldsymbol{F} = (\boldsymbol{m} \cdot \nabla)\boldsymbol{B}$$

where $\boldsymbol{m}$ is the amount of magnetisation each bead carries, the value of which is $8.4\times10^{-14}$ $Am^2$. Assuming the B field in the centre of the trap is 1 mT – a typical value during the tether experiments – and taking from Supplementary Fig. 1 $dx = 40\ \mu\text{m}$ and $dB = 0.002\% \times 1\ \text{mT}$, we get:

$$F = m\frac{dB}{dx} = 4.2 \times 10^{-5}\text{pN}$$

The magnetic force is 5 orders of magnitude lower than typical forces applied with the optical tweezers in the tether experiments.

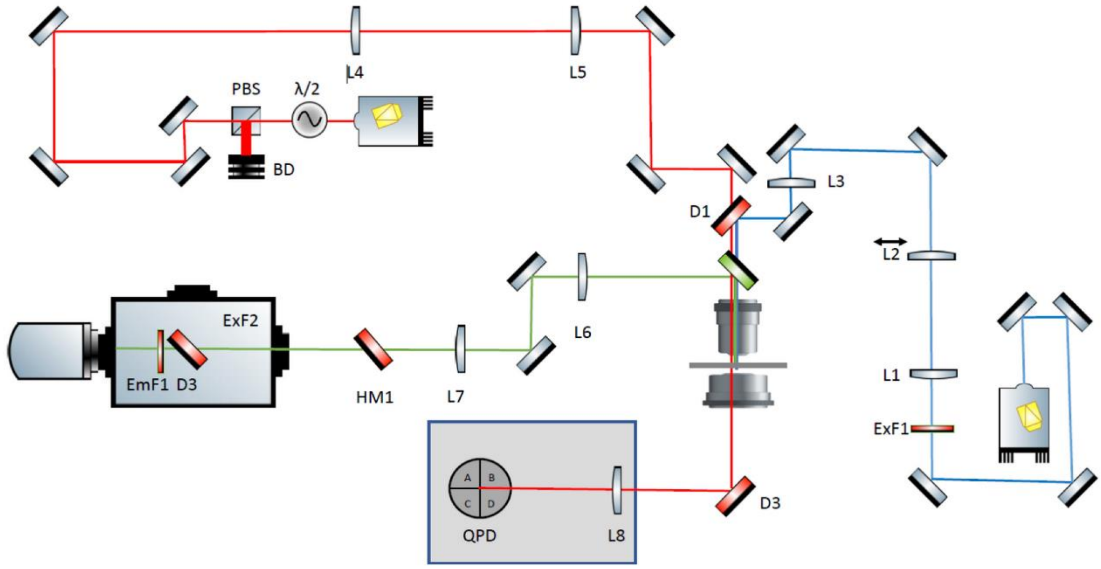


**Supplementary Figure 2**. **Schematics of the optical setup.** The 488 nm wavelength excitation laser beam is expanded 3x with the L1 and L2 lens pair, of which L2 is mounted on a horizontal translational lens mount for oblique illumination. The beam is then focused on the objective back-aperture by L3. The near-IR beam first goes through a quarter-wave plate mounted on a Thorlabs continuous rotation mount, allowing the subsequent polarized beam splitter (PBS) to redirect a controlled fraction of the beam onto a beam dump (BD). The remaining beam is then expanded 4x by a pair of lenses L4 and L5 before entering the back-aperture of the objective and overfilling it. The fluorescence signal is imaged with an Andor Ixon Ultra camera. The optical trap light is collected by an oil immersion condenser, redirected, and focused on a quadrant photodiode for force and displacement measurement of the bead.

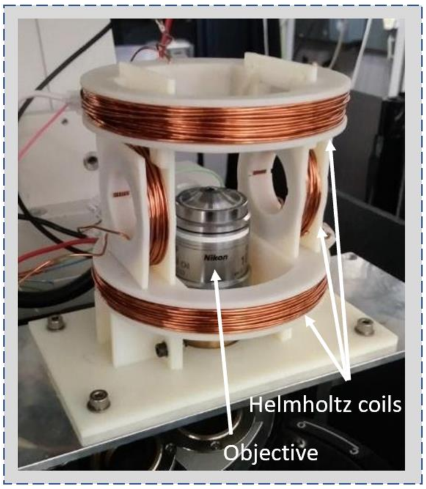


**Supplementary Figure 3. The Helmholtz coil pairs.** The photograph features the Helmholtz coil pairs and shows their positioning around the objective. The spools are 3D printed and the enamelled copper wires hand wound into the grooves of the spools. The nanostage is removed and the condenser arm is raised for clearer view.

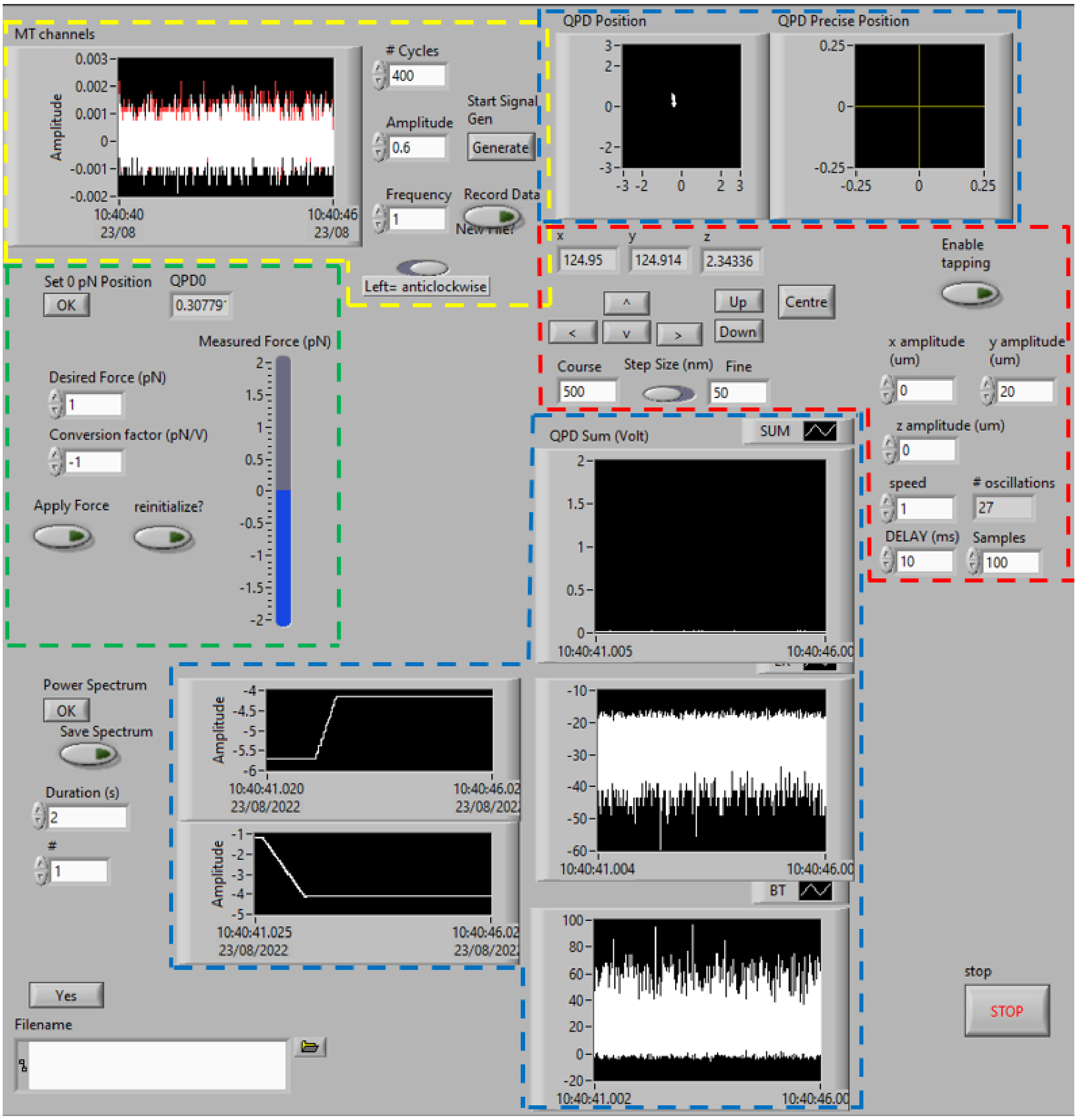


**Supplementary Figure 4**. **Screenshot of the microscope control Labview code GUI (Labview 2019).** The area inside of the red dashed lines is used for nanostage control and allows stepwise, continuous, and oscillating movement in 3D. The area inside of yellow dashed lines is used for MT control and real-time display of the current running through each coil. The areas inside of the blue dashed lines are used for real-time display of the voltages outputted by the QPD and nanostage. The area inside of the green dashed lines is used to set up and activate force-clamping. The software allows timestamped data acquisition across 12 different channels and signal generation across 10 at 80 kHz through an NI DAQ system (3 NI 9222 cards and 1 NI 9263, card, mounted on a NI cDAQ 9174, connected to the computer through USB).

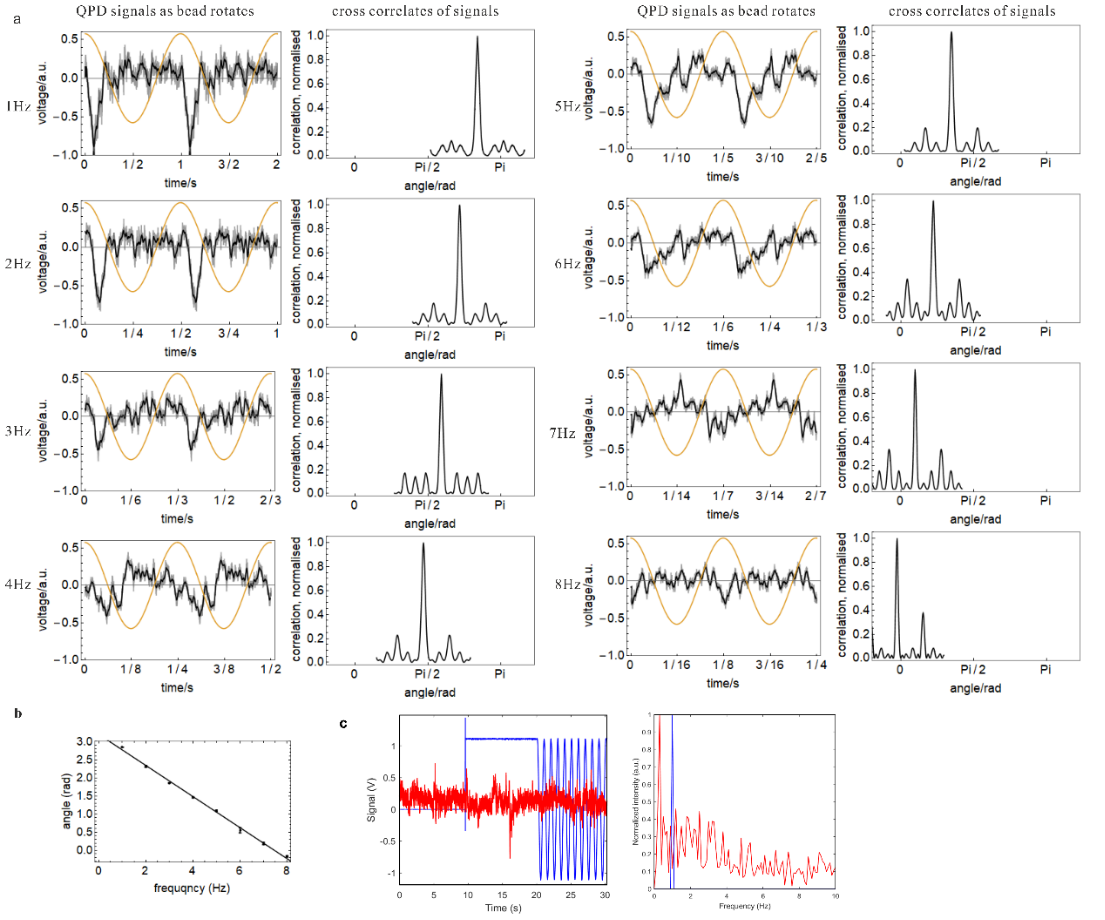


**Supplementary Figure 5. Calibration of the magnetic tweezers.** The magnetic field is set to rotate at increasing speed from 1 to 8 revolutions per second, which results in increasing drag that the bead experiences in the flow-cell buffer. This drag manifests as an angular lag between the angle of the B field and the angle of the bead's preferential axis, which is then used to calibrate the rotational stiffness of the magnetic tweezers. (a) The control voltage applied to the smaller pair of coils plotted on top of the QPD signal with its corresponding low-pass signal (signals above 10 Hz removed). The cross-correlation of the raw signals with the reverse of themselves are used to find the phase difference between adjacent frequencies. The peaks are marked to compare phases of the plots from different rotational frequencies. (b) The stiffness of the trap for the particular bead used to create the above plot is $1.1 \times 10^3$ pN·nm·rad$^{-1}$. The error bars show the standard errors from three measurements. (c) (left panel) QPD signal (red) is recorded initially with the B-field signal (blue) switched off, then switched on at 1 Hz after 10 s, (right panel) equivalent power spectrum indicating no influence of the rotating field on the QPD detection.

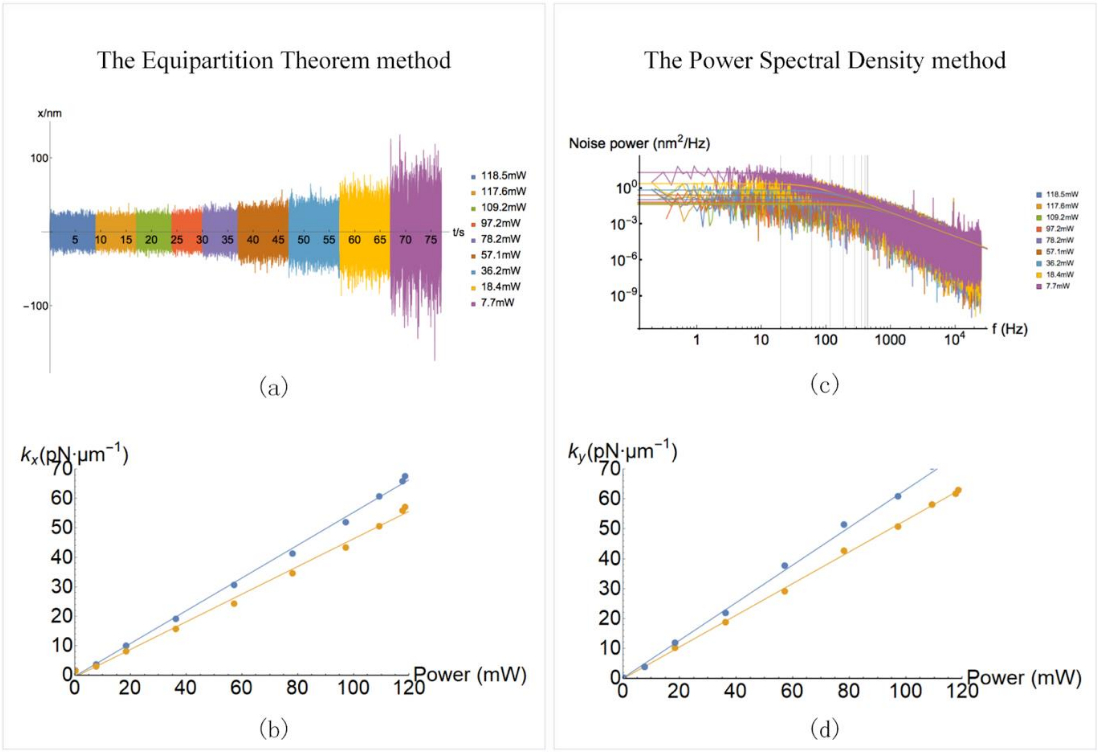


**Supplementary Figure 6. Calibration of the optical tweezers.** This is carried out both with the equipartition theorem method and the power spectrum method. a) Displacement trace of a trapped bead at various laser powers; b) Stiffness of the trap in the x- and y-directions as function of laser powers, obtained with the equipartition theorem method; c) PSD at various laser powers; d) Stiffness of the trap obtained with the power spectrum method.

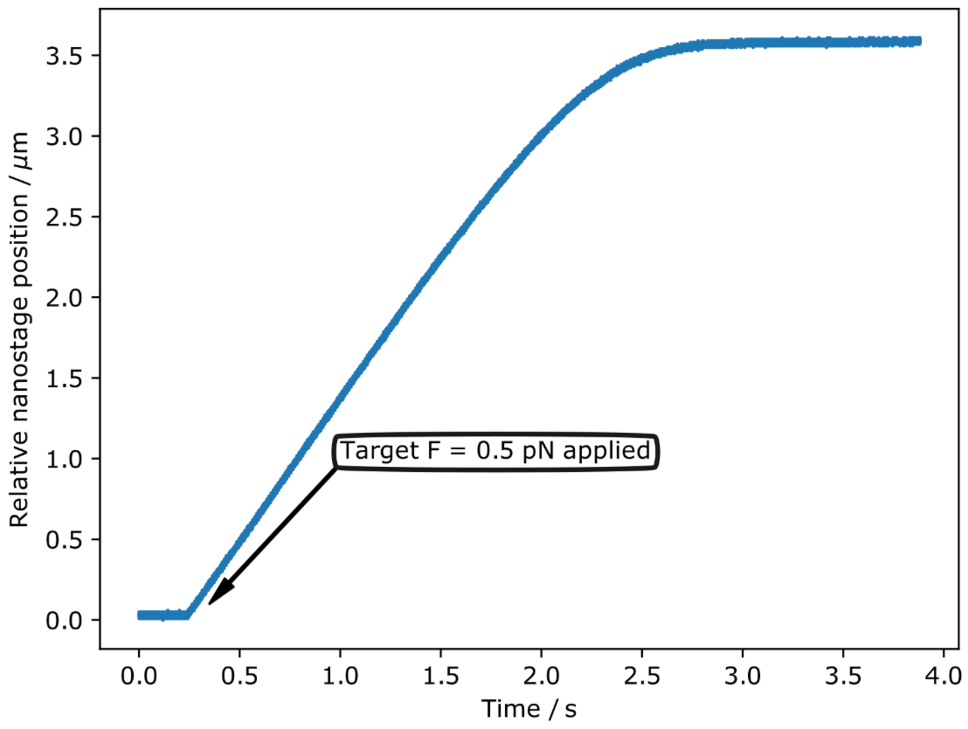


**Supplementary Figure 7. Force feedback response time is ~1 s.** Response of force clamp to instaneously applied force, reaching ~50% of required nanaostage displacement for the set force level here of 0.5 pN from a zero start point after approximately 1 s following the force perturbation.

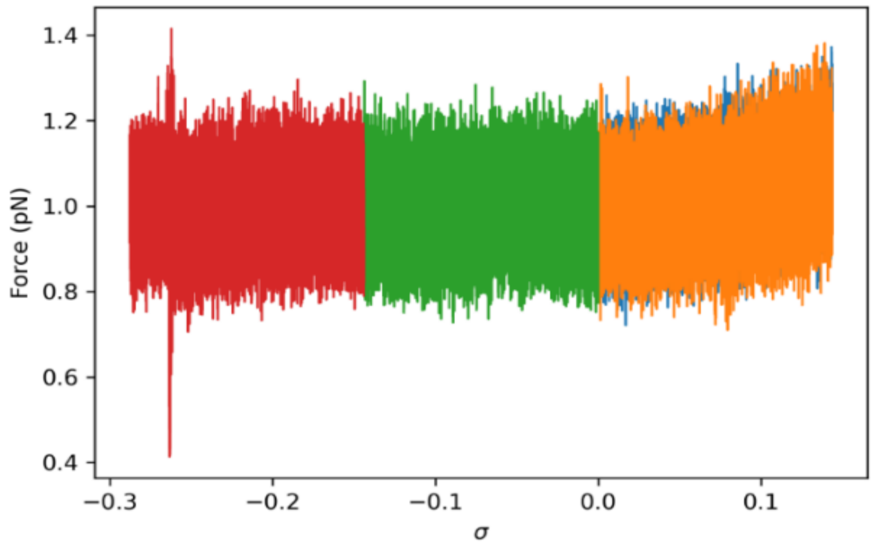


**Supplementary Figure 8**. **Set force using force clamp remains constant ±0.1 pN.** Clamping remains throughout an over- and undertwisting experiment, here shown at a set force of 1 pN displayed on the practical working force scale of 0-5 pN of COMBI-Tweez, root mean square noise ~0.1 pN.

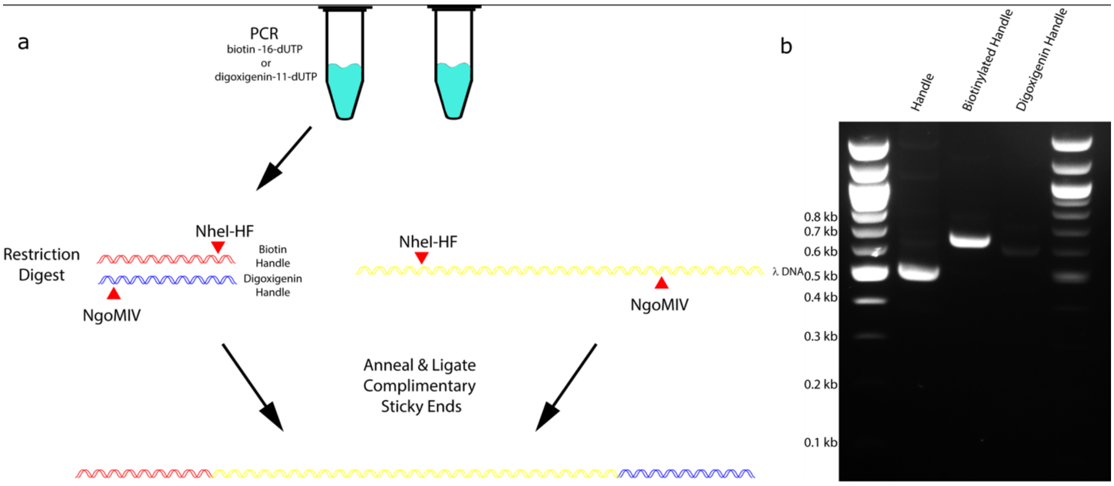


**Supplementary Figure 9. Schematics of the DNA tether construction protocol.** a) Diagrammatic representation for the creation of a DNA tether based on λ DNA and the pBS(KS+). Note the products created using the same primers but incorporating biotin-16-dUTP or digoxigenin-11-dUTP are retarded in the gel electrophoresis in comparison to the PCR product in the absence of these modified nucleotides. This suggests that these products do contain biotin or digoxigenin moieties.

### Supplementary Note 2: Steady-State Temperature Profile around an Isothermal Bead

#### I. INTRODUCTION

We are interested in the temperature profile around beads in an optical trap. In the following, we will first investigate how the distribution of magnetite in the bead may affect the heat absorbed. Secondly, we will assume spherical symmetry and solve the heat equation to calculate the temperature profile. We then conclude how the structure of the bead and the design of the optical trap may be engineered to minimise heating effects.

#### II. POWER ABSORPTION OF BEADS

We are interested in how the absorbed power decreases as the magnetite in the particle is distributed towards the surface rather than the centre of the bead. In general, the total amount of absorbed power can then be measured as

$$Q = \int \mathrm{d}\rho \alpha(\mathbf{r}) P(\mathbf{r}), \tag{1}$$

with $P$ the power distribution and $\alpha$ the local absorption coefficient of the material in the bead. We will conduct our studies by investigating a spherically symmetric bead ($\alpha(\mathbf{r}) = \alpha(\rho)$, with $\rho$ the distance from the centre of mass), in a uniform power distribution ($P(\mathbf{r}) = \mathrm{constant}$) and a uniform cylindrical beam ($P(\rho) = P(r)$, with $r$ the distance from the $z$ axis).

In this work, we create spherically symmetric beads by uniformly distributing the magnetite in a spherical shell between a radius $R_1$ and $R_2$ Thus, we move magnetite towards the surface by increasing $R_1$, and change $R_2$ such that the total volume of magnetite

$$V_\mathrm{m} = \frac{4\pi}{3}(R_2^3 - R_1^3). \tag{2}$$

remains constant. In a uniform power distribution all magnetite is uniformly heated, thus

$$Q = P\alpha V_\mathrm{m}, \tag{3}$$

independent of the choice of $R_1$.

In the following, we will investigate how the absorbed power will decay depending on the distribution of magnetite in the bead and on the power distribution of an optical trap.

##### A. Toy Model: Uniform cylindrical power distribution

We first developed a qualitative model to describe the power distribution in the laser beam as a simple cylinder of light. The $R_1$ dependence enters if a cylindrical beam has a smaller radius than the bead. The total volume hit by the beam with radius $R_\mathrm{beam}$ is composed of a cylinder with radius $\pi R_\mathrm{beam}$ and height $2\sqrt{R_2^2 - R_\mathrm{beam}^2}$, and two spherical caps with height $R_2 - \sqrt{R_2^2 - R_\mathrm{beam}^2}$. Combined, this gives

$$V(R; R_\mathrm{beam}) = \frac{4}{3}\pi R^3 \begin{cases} 1 - \left(1 - [R_\mathrm{beam}/R]^2\right)^{3/2}, \text{ for } R_\mathrm{beam} < R \\ 1, \text{ for } R_\mathrm{beam} \geq R. \end{cases}, \tag{4}$$

where $R = R_2$. Only the part of this volume that contains magnetite is heated; the part of the volume that is not heated is also composed of a cylinder with two spherical caps, but this time with $R = R_1$. Thus, the total volume of material that absorbs the uniform cylindrical beam is $V(R_2; R_\mathrm{beam}) - V(R_1; R_\mathrm{beam}) \equiv V_2 - V_1$. This gives for the absorbed power

$$Q = P\alpha(V_2 - V_1). \tag{5}$$

If the outer radius of the spherical shell, $R_2$, is smaller than the radius of the bead, all of the magnetite is heated uniformly, $V_2 - V_1 = V_\mathrm{m}$ and we retrieve Eq. (3). If the inner radius of the spherical shell is larger than the beam, we get

$$V_2 - V_1 = V_\mathrm{m} - \frac{4\pi}{3}\left[R_2^3\left(1-\left(1-[R_\mathrm{beam}/R_2]^2\right)^{3/2}\right) - R_1^3\left(1-\left(1-[R_\mathrm{beam}/R_1]^2\right)^{3/2}\right)\right]. \tag{6}$$

In the limit of large $R_1$, this reduces to

$$V_2 - V_1 = \frac{V_\mathrm{m}}{4\pi R_1^2}, \tag{7}$$

which shows that the amount of heated material decreases as $R_1^{-2}$ as the magnetite is distributed more towards the surface of the beads, provided that the beads are very large compared to the beam.

##### B. Power Distribution of Optical Trap

To make more robust quantitative comparisons with the experimental data, we then developed a more physical model for the power profile of the optical trap, which is given by

$$P(r,z) = P_0 \exp\left(-\frac{2r^2}{\omega_0^2}\left[1+\left(\frac{\lambda z}{\pi\omega_0^2}\right)^2\right]^{-1}\right), \tag{8}$$

where $\omega_0$ measures the width of the trap near the focus, and $\lambda$ describes the broadening of the trap away from the focus in both $z$ directions.

In the absence of profile broadening, the profile is spherically symmetric, and the absorbed power is given by

$$P = 4\pi\alpha P_0 \int_{R_1}^{R2} r^2 \tag{9}$$

###### 1. *Spherically symmetric Gaussian trap (small $\lambda$ limit)*

For small values of $\lambda$ the trap becomes spherically symmetric, and we can calculate the absorbed power as

$$Q = 4\pi\alpha P_0 \int_{R_1}^{R_2} r^2 \exp\left(-\frac{2r^2}{\omega_0^2}\right), \tag{10}$$

which can be solved analytically. We here just focus on the asymptotic limits where the amount of magnetite is either very small or very large. For the case where the volume of magnetite, $V_\mathrm{m}$, is very low, we find

$$\lim_{V_\mathrm{m}\to 0} Q = \frac{4\pi}{3}\alpha P_0 V_\mathrm{m} \exp(-2R_1^2/\omega_0^2), \tag{11}$$

which is a Gaussian with width $\omega_0/2$. This width remains within the same order of magnitude in the limit where the volume of magnetite is very large. In this case we have

$$\lim_{V_\mathrm{m}\to\infty} Q = \frac{\pi}{4}\alpha P_0\left(\sqrt{2\pi} + 4\frac{R_1}{\omega_0}\exp(-2R_1^2/\omega_0^2) - \sqrt{2\pi}\,\mathrm{erf}(\sqrt{2}R_1/\omega_0)\right), \tag{12}$$

which indicates the absorbed power decays with a semi-Gaussian profile as $R_1$ increases. The most rapid decay occurs at the inflection point, $R_1 = \omega_0/\sqrt{2}$.

##### 2. Axial dependence

We formulate the integral in spherical coordinates (using $z = r\cos\theta$) as

$$Q = 2\pi\alpha P_0 \int_{R_1}^{R_2} \mathrm{d}r \int_0^{\pi} \mathrm{d}\theta\, r^2 \sin\theta \exp\left(-\frac{2r^2}{\omega_0^2}\left[1+\left(\frac{\lambda r\cos\theta}{\pi\omega_0^2}\right)^2\right]^{-1}\right). \tag{13}$$

This integral can be solved numerically (e.g. using Mathematica's `NIntegrate` function)

In the limit where $R_1$ is large compared to the spherically symmetric component ($R_1 \gg \omega_0$), this equation can be solved analytically, and gives

$$\lim_{R_1\to\infty} Q = \alpha P_0 V_\mathrm{m}\left(\exp\left(-\frac{2\pi^2\omega_0^2}{\lambda^2}\right) - \frac{2\omega_0}{\lambda}\pi^{3/2}\mathrm{erfc}\left(\sqrt{2}\pi\omega_0/\lambda\right)\right). \tag{14}$$

If $\lambda \gg \omega_0$, then this reduces to $Q = \alpha P_0 V_\mathrm{m}$, indicating all supplied power may be absorbed by magnetite (thus, limited only by the absorption coefficient $\alpha$). In the opposite case $\lambda \ll \omega_0$ no power is absorbed, as expected from the spherically symmetric case. The axial component has a negligible contribution to power absorption for small values of $\lambda$; for $\lambda < 2.8\omega_0$ the fraction of magnetite involved in power absorption is less than $0.01$. However, at the inflection point at $\lambda \approx 6.7864\omega_0$ approximately a quarter of the magnetite is involved in power absorption, while half of the magnetite becomes involved at $\lambda \approx 12.7038\omega_0$.

#### III. TEMPERATURE PROFILE: SPHERICAL SYMMETRY

We consider a spherical bead with a radius $R_0$ with a fixed temperature $T_0$ in stagnant water with a bulk temperature $T_\mathrm{room}$, and we are interested in the steady-state temperature profile $T(r)$ as a function of the distance $r \geq R_0$ from the centre of mass. At large distances, heat transfer is dominated by free convection while at short distances conduction is the dominant mode of heat transfer. The crossover distance, $L$, between these regimes can be calculated using the Nusselt number

$$\mathrm{Nu} = \frac{hL}{k}, \tag{15}$$

which equals $2$ for a sphere in a stagnant medium. For water, the conductive heat transfer coefficient is $k = 0.6$ W/m·K, and the convective heat transfer coefficient is $h = 500 - 10{,}000$ W/m$^2$ K. Hence, the characteristic length beyond which convection becomes important is $L = 10 - 500\,\mu$m. Thus, in our experiments the relevant length scales are relatively short and heat transfer is dominated by conduction. Under these conditions, heat transfer is governed by the heat equation

$$\rho C_\mathrm{p}\frac{\partial T}{\partial t} = k\nabla^2 T + q, \tag{16}$$

where the Laplacian is in spherical coordinates given by

$$\nabla^2 T = \frac{1}{r^2}\frac{\partial}{\partial r}\left(r^2\frac{\partial T}{\partial r}\right), \tag{17}$$

and where $\rho$ is the density, $C_\mathrm{p}$ is the heat capacity, and $k$ is the conductive heat transfer coefficient, and which are all materials dependent properties. The source term $q(r)$ depends on the power distribution of the light source and on the light absorption coefficient of the material.

In the following, we will consider a spherical bead composed of a polystyrene core for $r < R_1$, a magnetite shell for $R_1 \leq r \leq R_2$, and a polystyrene shell for $R_2 \leq r \leq R_\mathrm{bead}$, and which is surrounded by water for $r > R_\mathrm{bead}$. Heat is generated in the magnetite shell ($q > 0$), and dissipates through heat transport through the polystyrene shell and water. We will assume steady state conditions. This ensures that the temperature in the polystyrene core is homogeneous, while the temperature profile in the magnetite shell is governed by

$$k_\mathrm{m}\nabla^2 T + q = 0, \text{ for } R_1 < r \leq R_2 \tag{18}$$

in polystyrene by

$$k_\mathrm{PS}\nabla^2 T = 0, \text{ for } R_2 < r \leq R_\mathrm{bead} \tag{19}$$

and in water by

$$k_{\rm water}\nabla^2 T = 0,\ \text{for } R_{\rm bead} < r < \infty. \tag{20}$$

From these equations, we find that the total heat generated is

$$Q = \int_{R_2}^{R_3} \mathrm{d}r q(r). \tag{21}$$

For a uniform power distribution,

$$Q = \frac{4}{3}\pi(R_2^3 - R_1^3)q, \tag{22}$$

In steady state, this uptake of energy must be balanced by the dissipation of energy through heat conduction. Due to spherical symmetry, this implies that the flow rate of heat is constant, thus

$$4\pi r^2 k\nabla T = Q, \tag{23}$$

with $\nabla T = \partial T/\partial r$. This condition, the room temperature at large distances $r$, and continuity of the temperature at the interfaces (assuming efficient heat transfer across the interfaces), will set the boundary conditions of differential equations defined above. Below we will solve them, starting from the bulk water, followed by an evaluation of the polystyrene and the magnetite shells, respectively.

##### A. Temperature in the water outside the bead

In water, the temperature profile is set by $\nabla^2 T = 0$. In spherical coordinates, this gives the general solution

$$T(r) = A + B\frac{1}{r}. \tag{24}$$

Far away from the bead (so for large $r$), this must decay to room temperature, thus $A = T_{\rm room}$. The heat flux is $4\pi r^2 k_{\rm water}\nabla T = 4\pi B$, and must balance the heat generation in Eq. 22. This gives $B = (R_2^3 - R_1^3)q/3k_{\rm water}$, and yield the temperature profile in the water phase,

$$T(r) = T_{\rm room} + \frac{Q}{4\pi k_{\rm water}}\frac{1}{r},\ \text{for } r \geq R_{\rm bead}, \tag{25}$$

which may alternatively be written as

$$\boxed{T(r) = T_{\rm room} + (T_{\rm surface} - T_{\rm room})\frac{R_{\rm bead}}{r},\ \text{for } r \geq R_{\rm bead},} \tag{26}$$

with the surface temperature

$$T_{\rm surface} = T_{\rm room} + \frac{Q}{4\pi k_{\rm water}}\frac{1}{R_{\rm bead}}. \tag{27}$$

This equation is independent of how the source of heat is distributed within the bead, and applies as long as this source is spherically symmetric.

##### B. Temperature in the polystyrene shell

In the polystyrene shell, we again have the general solution

$$T(r) = A + B\frac{1}{r}, \tag{28}$$

where as before $B = Q/4\pi k_{\rm PS}$ ensures that the dissipated heat matches the generated heat. The coefficient $A$ is set by the surface temperature, $T_{\rm surface} = A + Q/4\pi k_{\rm PS}R_{\rm bead}$, and so we have

$$T(r) = T_{\rm surface} + \frac{Q}{4\pi k_{\rm PS}}\left(\frac{1}{r} - \frac{1}{R_{\rm bead}}\right),\ \text{for } R_2 \leq r \leq R_{\rm bead}, \tag{29}$$

which gives

$$T_2 = T_{\rm surface} + \frac{Q}{4\pi k_{\rm PS}}\left(\frac{1}{R_2} - \frac{1}{R_{\rm bead}}\right). \tag{30}$$

as the interfacial temperature between magnetite and polystyrene at $r = R_2$.

##### C. Temperature in the magnetite shell

In the magnetite shell, the temperature profile is described by $k_{\rm m}\nabla^2 T + q = 0$, which has the general solution

$$T(r) = \frac{q}{6k_{\rm m}}r^2 + A + B\frac{1}{r}. \tag{31}$$

At $r = R_2$, the heat flux $4\pi R_2^2 k_{\rm m}\nabla T$ must match the total generated heat $Q$ to ensure a steady state. This results in $B = Q/4\pi - qR_2^3/3k_{\rm m} = -qR_1^3/3k_{\rm m}$. To obtain $A$, $T(R_2)$ must match the interfacial temperature $T_2$, thus $A = T_2 - qR_2^2/6k_{\rm m} - B/R_2$. This gives for the steady-state temperature profile in magnetite

$$T(r) = T_2 - \frac{q}{6k_{\rm m}}(r^2 - R_2^2) - \frac{qR_1^3}{3k_{\rm m}}\left(\frac{1}{r} - \frac{1}{R_2}\right). \tag{32}$$

#### IV. RESULTS

Experimentally, the surface temperature can be determined by bringing the bead into contact with a wax with melting temperature $T_{\rm m}$ and determining the distance $R_{\rm m}$ at which the wax melts. Using Eq. (26) the surface temperature can then be calculated as

$$T_{\rm surface} = T_{\rm room} + (T_{\rm m} - T_{\rm room})\frac{R_{\rm m}}{R_{\rm bead}}. \tag{33}$$

We have determined $R_{\rm m}$ for three types of wax with different melting temperatures. The distance-dependence of the temperature does not obey the expected $1/r$ dependence. This may be due to heating the solvent, or due to the coverslip that may act as a heatsink. For the latter reason, we may only expect the $1/r$ dependence to hold for a measured $R_{\rm m}$ smaller than approximately $2.5$ $\mu$m (the radius of the bead is 1.5$\mu$m, and the distance of the bead from the coverslip is 1 $\mu$m). Thus, only the wax with $T_{\rm m} = 42°$C could be used, which melted at a distance $1.723 \pm 0.389$ $\mu$m from the centre of the bead, giving an estimate of $45.1 \pm 5.4°$C.

#### V. CONCLUSION

We have discussed several factors that affect the temperature profile around the bead. First, we have discussed how the absorbed power is determined by the size of the bead and the distribution of material within the bead. The absorbed energy is maximised if magnetite is centered in the core of the bead. It can be considered maximally centered if all material is within the focus of the optical trap, which is set by the length scale $\omega_0$. As the magnetite is distributed further from the focus, the absorbed power decays according to a Gaussian tail. The power vanishes to negligible values if $\lambda < 3\omega_3$, where $\lambda$ describes the $z$ component of the optical trap. If $\lambda$ increases, then the amount of absorbed energy becomes finite; As $\lambda \approx 6.78\omega_0$, the absorption of magnetite would remain $24\%$ as strong as if it wear in the centre of the bead; for $\lambda \approx 12.7\omega_0$ this increases to $50\%$. Thus, the amount of absorbed energy can be engineered through both the design of the magnetic bead and by the properties of the optical trap. Assuming the surface temperature of the bead is uniform, the temperature profile in the water near the bead is expected to decay via thermal conduction (rather than convection) via a $1/r$ dependence. This dependence follows from spherical symmetry arguments, and breaks down at the larger distances where heat transport is altered by the coverslip.

**Supplementary Note 3: Finite element modelling of laser heating of the optical trap with a magnetic bead**

**Heat transport.** Sample heating can be described with the heat equation. Since in only 1 ms the temperature reaches 90% of the steady-state value[1], we neglect the time-dependant term. At equilibrium, the heat equation is:

$$\nabla^2 T = -\frac{q}{K} \tag{34}$$

where $T$ is the temperature, $K$ is the thermal conductivity and $q$ is the energy absorbed per unit volume per unit time. Eq. 34 describes how energy transferred to the bead diffuses throughout the bead and the surrounding buffer solution. Since the magnetic bead is made of polystyrene with a non-uniform distribution of magnetite nanoparticles embedded inside, we need to consider the energy absorbed per unit volume per unit time separately for regions of magnetite and regions of polystyrene.

**Magnetic bead structure.** TEM data indicates that the magnetic bead has a core-mantle-crust structure (main text Fig. 2a-c). The core has a radius of 1.35 ± 0.05 µm and, indicated by the Micromer-M bead manufacturer, it is made of styrene-maleic acid copolymer. The core is wrapped in a layer of magnetite whose thickness is approximately 100 nm. The crust is a thin layer of styrene-maleic acid copolymer less than 100 nm in thickness.

**3D laser profile.** The laser beam has a Gaussian profile at the focal plane, with irradiance:

$$I(r,z) = I_0 \exp(-\frac{2(r/r_0)^2}{\omega(z/z_0)^2}) \tag{35}$$

where $r$ is the radius, $I_0$ is the peak irradiance at the centre of the beam, $r_0$ and $z_0$ are constants, and $z$ is the axial coordinate, $\omega(z/z_0)$ is the radius of the laser beam where the irradiance is $1/e^2$ (13.5%) of the maximum irradiance. The radius of the beam is:

$$\omega(z) = \omega_0 \sqrt{1 + (\frac{\lambda z}{\pi \, \omega_0{}^2})^2} \tag{36}$$

where the wavelength λ is 1064 nm, $\omega_0$ is the radius of the beam at the waist. We take the approximation that $\omega_0 \approx \lambda / (NA \cdot \pi)$ where $NA$ is the numerical aperture of the objective lens.

Due to the axial optical force and the relatively high opacity of the magnetite, the trapped bead is positioned $\sim 2$ µm above the centre of the trap. Also, the bead undergoes Brownian motion, so from the perspective of the bead, the laser profile will be spread out along both the *r*- and *z*-axis. Thus, the modification factors $r_0$ and $z_0$ are added in the expression of $I(r,z)$. The root mean square of the Brownian motion is a few tens of nanometres.

**Energy absorption.** We take the above core-mantle-crust structure of the bead and the irradiance distribution of the laser to obtain the energy absorbed per unit of volume and unit of time:

$$q(r,\theta,\varphi) = \alpha(r) \cdot I(r,z) \cdot e^{-\alpha(r)} \tag{37}$$

where $\alpha(r)$ is the absorption coefficient, which depends on the materials, i.e. 10 $m^{-1}$ for $0 < r <$ 1.3 µm (polystyrene), 2000 $m^{-1}$ for $1.3 < r < 1.4$ µm (magnetite)[2] and 10 $m^{-1}$ for $1.4 < r < 1.5$ µm (polystyrene). The last term $e^{-\alpha(r)}$ considers the attenuation of the laser irradiance as the light penetrates the layers of materials in the bead.

**Simulation of energy absorption.** The NIR laser power was nominally set to 100 mW. We centred the bead at $z = 0$ and $r = 0$ so the centre of the laser trap will be at $z = -2$ μm; the numerical aperture of the objective *NA* = 1.49; the modification factors to consider Brownian motion are taken to be $r_0 = 0.90$ and $z_0 = 0.95$ respectively. We used MATLAB to numerically evaluate Eq. (37) for a 3D mesh of positions throughout the volume of the magnetic bead. Then we integrate *q* over the polystyrene core, the magnetite mantle and the magnetite crust separately to obtain the heat generation of the individual layers.

**Temperature simulation.** The thermal conductivity of polystyrene is $K_p$ = 0.12 W/mK, that of magnetite is $K_m$ = 4 W/mK, and that of water is 0.597 W/mK.

For the geometry, the coverslip surface is 1 μm away from the edge of the bead but the microscope slide is ignored so the other side is taken as an infinite water bath. The conductivity of the coverslip is $K_c$ = 1.4 W/mK; the temperature at the coverslip surface is assumed to be above the room temperature and the exact value of the temperature at the coverslip surface is obtained by fitting the simulated temperature curve to experimentally measured temperatures at several points in the surroundings of the bead; the temperature at $r = \infty$ for the water bath is set to 21°C.

We used the MathWorks Partial Differential Equation (PDE) Toolbox to numerically solve Eq. (34). For the magnetic bead, we used the function `multisphere` to create the three-layered geometry. It supports specification of thermal conductivities, *K*, and heat sources, *q*, for individual layers in the bead. Also, it manages the calculation of heat transfer at the contacting surfaces between layers.

Supplementary Fig. 10a,b show the results of the simulation. In panel (a), the temperature is sampled along a horizontal line through the centre of the bead. The temperature within the bead is relatively constant at slightly above 45°C. Away from the bead, the temperature initially falls quickly and then decreases slowly. The red data points indicate experimentally measured temperatures, which match the simulated trend well. In the horizontal plane through the centre of the bead, the temperature distribution is circularly symmetric. In the vertical plane (the corresponding contour plot not shown here), the temperature gradient between the bottom of the bead and the coverslip is large while that above the bead is small. This is due to the coverslip acting as a heat sink.

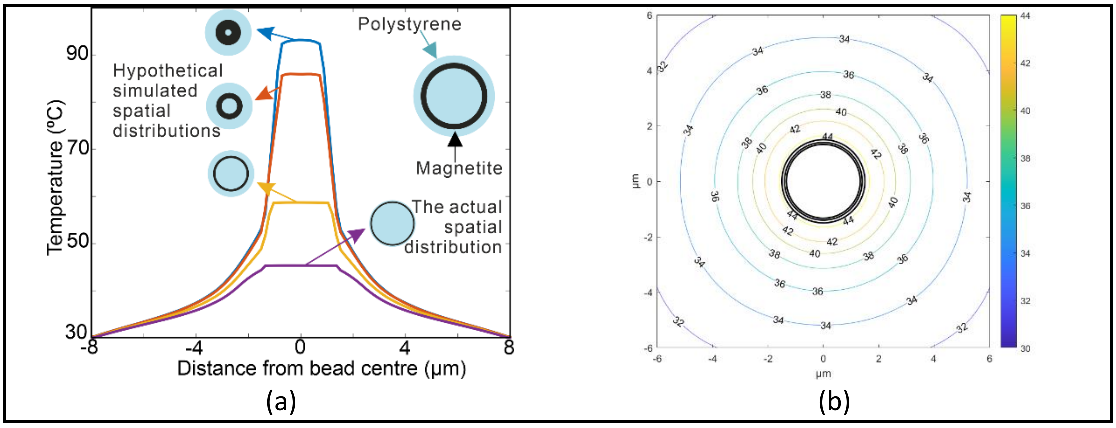


**Supplementary Figure 10.** (a) Finite element model for heat transfer predicts that having magnetite distributed closer to bead core generates much higher temperatures than the 45˚C we observe. (b) The temperature contour plots in the horizontal plane. The three black circles indicate the boundaries of the polystyrene core, the magnetite mantle and the magnetite crust of the bead.

It is worth noting that the exact surface temperature of the coverslip is unknown prior to the simulation although it is one of the input variables to the simulation. We optimised its value such that the simulated temperature curve crosses all three experimental points. This also allows us to obtain the surface temperature of the coverslip (26±1°C). The error indicates the range of values the temperature can take whilst keeping the simulated curve match the measured data. The same applies to the absorption coefficients of magnetite. Although a value of 2000 $m^{-1}$ is reported in literature[2], the configuration of their nanoparticles is not identical to ours so our absorption coefficients will likely be in the vicinity of but not exactly 2000 $m^{-1}$. We found it to be 4000±2000 $m^{-1}$.

We also simulated the magnetic bead configurations in which the magnetite nanoparticles are distributed closer to the centre of the bead. We postulate that when the magnetite nanoparticles are arranged closer to the centre, they will sit at regions of more intense laser radiation so they will absorb more heat and thus the temperature will be higher. Main text Fig. 2h shows the temperature simulation for various magnetite nanoparticle arrangements. In all plots, the total volume of the magnetite is kept the same. When the magnetite is in a thin shell less than 100 nm from the outer edge of the bead, the temperature at the surface of the bead is just above 45°C - this is the real-life configuration. When the magnetite is in a shell of inner radius 0.2 $\mu m$ and outer radius 0.82 $\mu m$, namely almost all centred in the middle, the temperature of the bead is above 90°C.

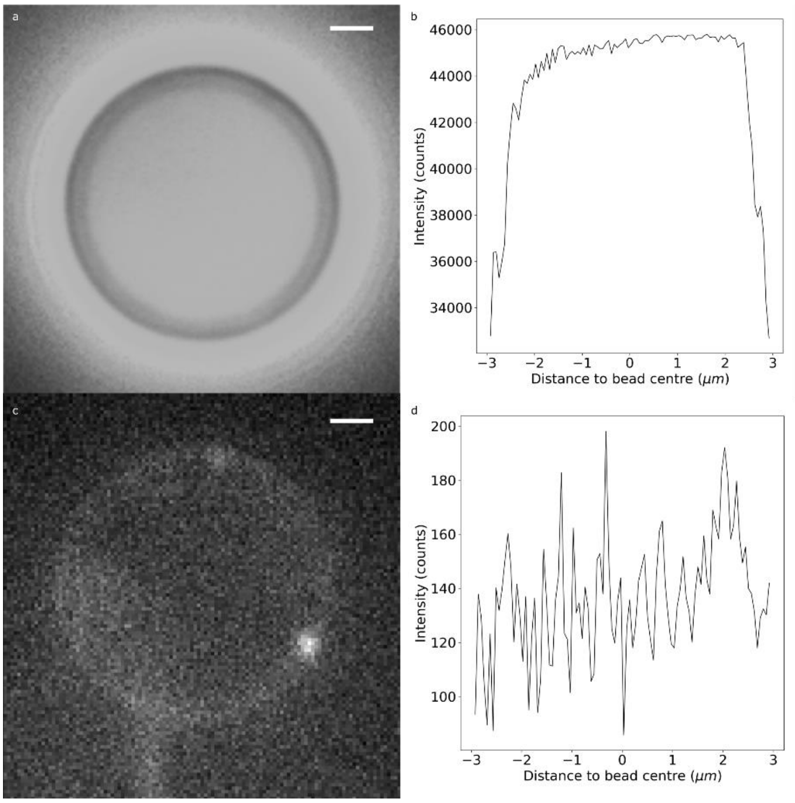


**Supplementary Figure 11. Characterisation of bead brightness in fluorescence.** Brightness of in-house functionalised anti-DIG beads (bottom) and commercial (Spherotech) beads (top) imaged under the same conditions. Bars: 1 µm.

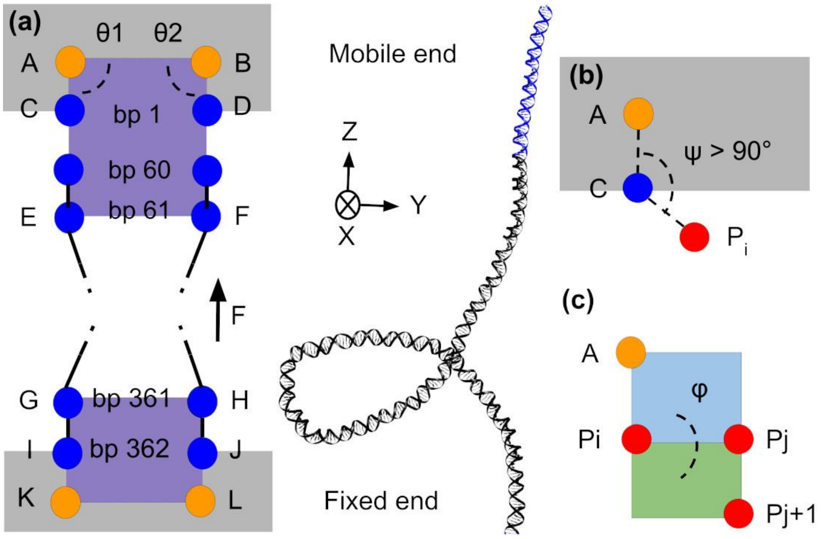


**Supplementary Figure 12**. **Schematic of the restraints applied in our simulations to control torsion and tension on linear DNA.** Dummy atoms, which act as a reference points, are pictured as orange circles. Key O3' or O5' atoms at each end are pictured as blue circles and phosphorus atoms are pictured as red circles. The motion of atoms C and D is limited to the Z axis, thanks to angular restrains such as $\theta_1$ and $\theta_2$. Grey areas represent excluded volume, in to which the bulk of the strand cannot move. Purple areas represent co-planar planes (ABE -BAF and KLH-LKG) responsible of maintaining the DNA torsionally constrained. The blue and green areas are the two planes that define the dihedral angle ϕ.

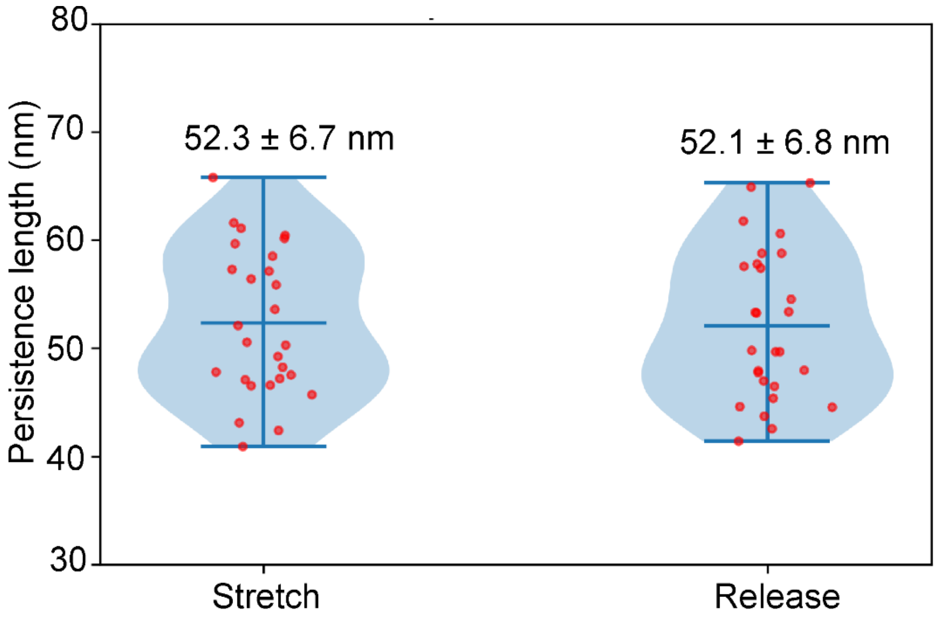


**Supplementary Figure 13. Stretch and release half cycles from the same DNA tether exhibit similar persistence lengths**. Violin plot showing distribution of fitted wormlike chain persistence lengths taken for the same DNA tether on separate consecutive stretch and release half cycles at 1Hz per whole cycle (fitted contour length 5.3 µm). Mean and s.d. indicated, n= 27 cycles, two-tailed Student's *t* test (making no assumption about distribution normality, no covariate analysis performed) P=0.9162 between stretch and release distributions (not significant).

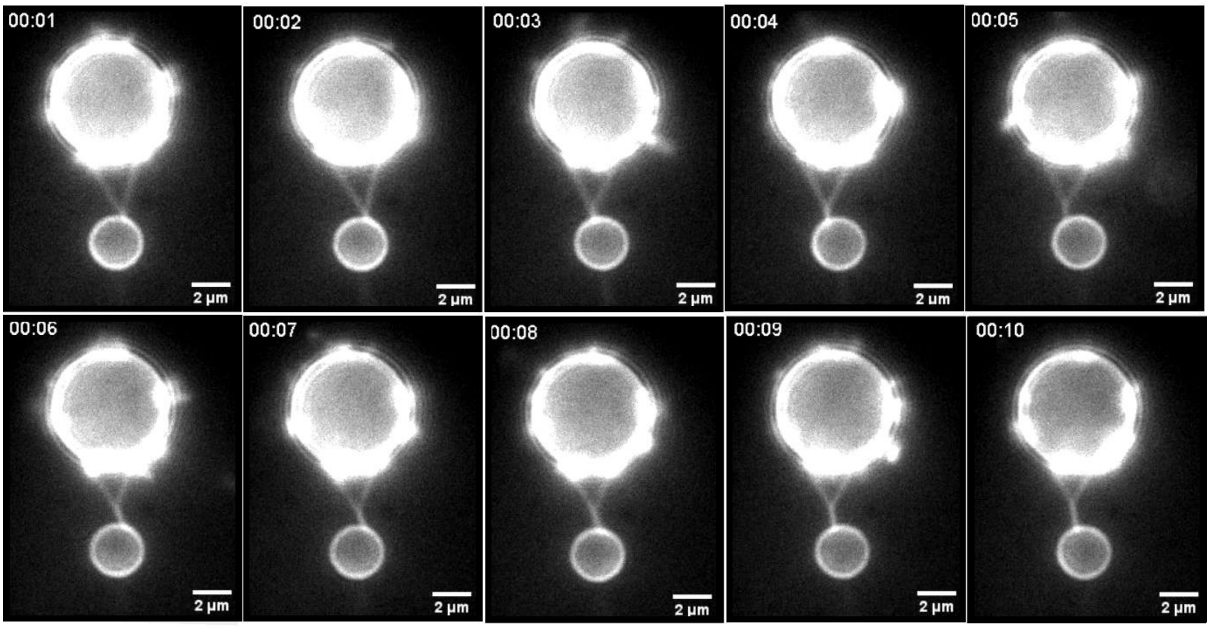


**Supplementary Figure 14. Controllably elevating the DNA surface density on a bead enables braiding to be explored.** Consecutive image frames from fluorescence microscopy (time stamp indicated in seconds) of a continuously overtwisted optically trapped magnetic bead (smaller bead, larger anchor bead immobilised to coverslip surface) rotated in the B-field of COMBI-Tweez at 1Hz, showing the braiding of two separate DNA tethers to form a growing braid interface.

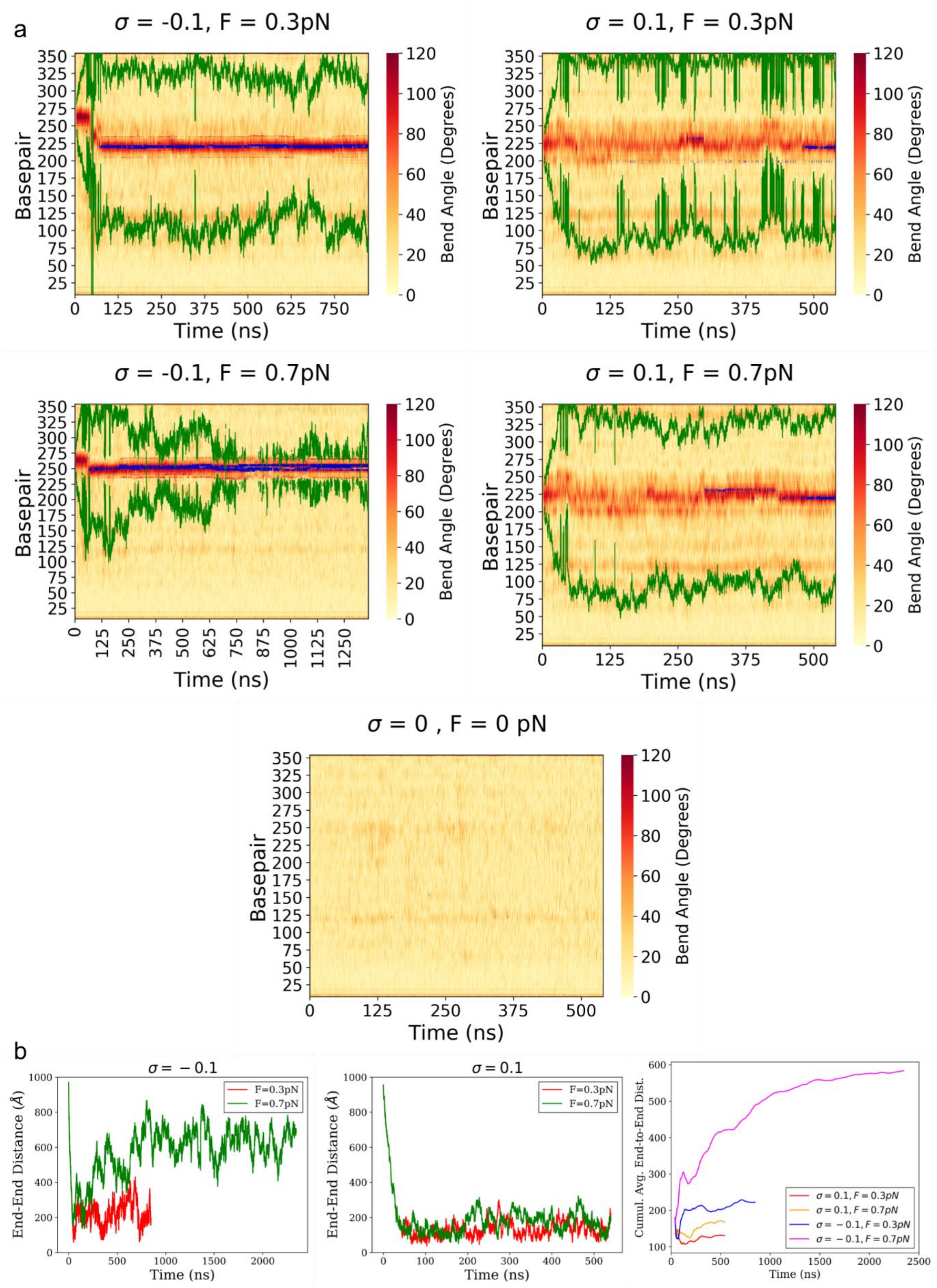


**Supplementary Figure 15.** a) Kymographs depicting dynamics of plectoneme boundaries (green lines) and denaturation bubbles (blue points) along with local (15 bp) bending angles (heatmap). Colocalisation is observed in all cases, although negatively supercoiled bubbles show significantly more stability than those in positive supercoiling. Three denaturation events were seen in relaxed

DNA (that is, at 0 pN and σ=0), but they do not meet our definition of melting bubbles because they are shorter than 1 ns. b) End-to-end distance and cumulative average of the end-to-end distance along time for our four simulations combining σ = ±0.1 and F = 0.3, 0.7 pN.

**Supplementary Movie Legends**

**Supplementary Movie 1**

Schematic video of the COMBI-Tweez system showing geometries alongside simulated and real data.

**Supplementary Movie 2**

Representative force extension experiment as performed in brightfield, see also Fig. 3. The maximum extension is set so that the optically trapped bead is just being pulled from the trap.

**Supplementary Movie 3**

An optically trapped bead tethered to an anchor bead is rotated, building up supercoiling density, forming plectonemes, and generating force. At a critical moment, the DNA buckles and the bead is pulled from the trap entirely. See also Fig. 3.

**Supplementary Movie 4**

DNA supercoiling with the force clamp applied. Here to keep the force constant the nanostage moves, reducing the distance between the anchor bead and optically trapped bead. See also Fig. 3.

**Supplementary Movie 5**

An anchor bead with single DNA molecules bound to the surface and imaged with SYBR Gold. To form a tether, a DNA molecule would be selected and the optically trapped bead brought into close proximity with it. See also Fig. 4.

**Supplementary Movie 6**

A force extension experiment performed during fluorescence imaging. See also Supplementary Movie 2 and Fig. 5.

**Supplementary Movie 7**

Two tethers between the anchor and trapped bead can be braided together by applying rotation to the optically trapped bead, increasing the braided region in the centre. See also Fig. 4.

**Supplementary Movie 8**

Two DNA molecules which are braided together are imaged, with one molecule breaking due to reactive oxygen species damage. The broken molecule retracts along the remaining tether. See also Fig. 4.

**Supplementary Movie 9**

A DNA tether imaged with SYBR Gold is seen breaking and retracting to the anchor bead due to entropic forces. See also Fig. 4.

**Supplementary Movie 10**

Video of a plectoneme formed by positive supercoiling. See also Fig. 5.

**Supplementary Movie 11**

Video of a DNA molecule buckling and pulling the optically trapped bead from the trap due to positive supercoiling.

**Supplementary Movie 12**

Video of a tethered full lambda DNA molecule following twisting show three plectonemes, which disappear sequentially following a single-strand DNA nick event followed by a torsional relaxation wave.

**Supplementary Movie 13**

MD simulation of the 362bp duplex at a supercoiling density of $\sigma = +0.1$ with a pulling force of 0.3 pN. See also Fig. 6.

**Supplementary Movie 14**

MD simulation of the 362bp duplex at a supercoiling density of $\sigma = +0.1$ with a pulling force of 0.7 pN. See also Fig. 6.

**Supplementary Movie 15**

MD simulation of the 362bp duplex at a supercoiling density of $\sigma = -0.1$ with a pulling force of 0.3 pN. See also Fig. 6.

**Supplementary Movie 16**

MD simulation of the 362bp duplex at a supercoiling density of $\sigma = -0.1$ with a pulling force of 0.7 pN. See also Fig. 6.

**Uncropped image of the gel shown in Supplementary Figure 9b:**

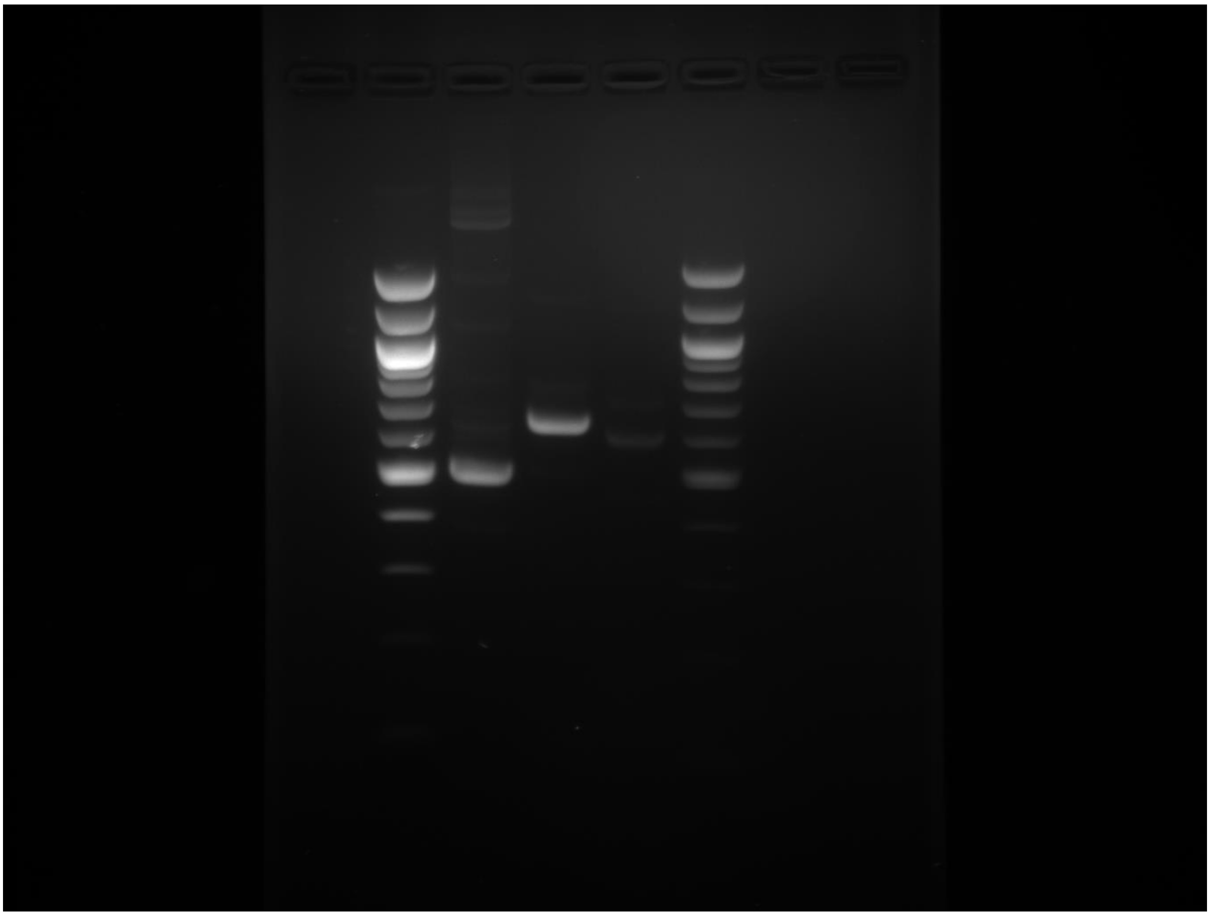
