## Supplementary figures and images for "Correlating fluorescence microscopy, optical and magnetic tweezers to study single chiral biopolymers such as DNA"

### Supplementary Movie 2

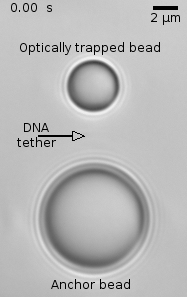

### Supplementary Movie 3

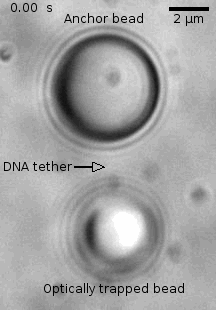

### Supplementary Movie 4

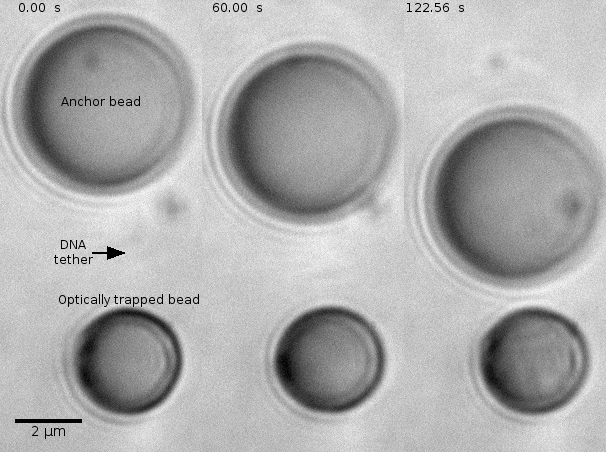

### Supplementary Movie 5

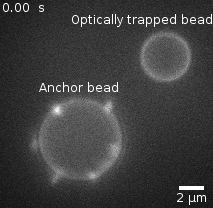

### Supplementary Movie 6

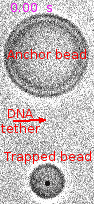

### Supplementary Movie 7

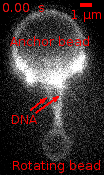

### Supplementary Movie 8

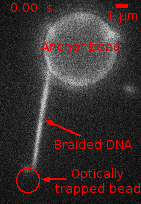

### Supplementary Movie 9

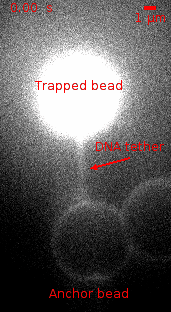

### Supplementary Movie 10

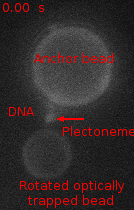

### Supplementary Movie 11

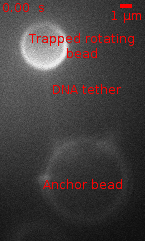

### Supplementary Movie 12

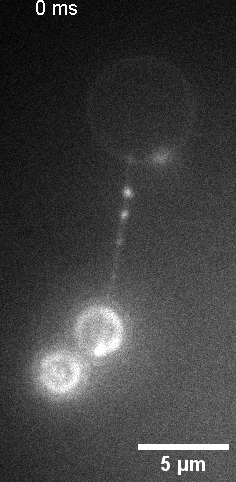
